# γ-Linolenic acid increases expression of key regulators of lipid metabolism in human skeletal muscle cells

**DOI:** 10.1101/605949

**Authors:** L.A Baker, N.R.W Martin, D.J Player, M.P Lewis

**Author notes:** Corresponding author: Dr. Luke Baker, Department of Health Sciences, University of Leicester, Leicester.

## Abstract

Control of skeletal muscle fat metabolism is regulated acutely through Peroxisome Proliferator Activated Receptor (PPAR) δ activation and its downstream intracellular targets. The purpose of this study was to determine whether fatty acids with high binding affinity for PPARδ can elevate the expression of genes related to fatty acid oxidation and indicators of mitochondrial biogenesis in cultured human skeletal myotubes. Myotubes were treated for 72 hours with one of four conditions: (i) Control (CON); (ii) Eicosapentaenoic acid (EPA 250μM); (iii) γ-linolenic acid (γ-LA 250μM); (iv) PPARδ Agonist (GW501516 10nM). Incubation with γ-LA induced increases in the gene expression of CD36 (p= 0.005), HADHA (p= 0.022) and PDK4 (p=0.025) in comparison with CON, with no further differences observed between conditions. Furthermore, intensity of MitoTracker® Red immunostaining in myotubes increased following incubation with γ-LA (p≤ 0.001) and EPA (p= 0.005) however these trends were not mirrored in the expression of PGC-1α as might be expected. Overall, γ-LA elevates levels the transcription of key intracellular regulators of lipid metabolism and transport in human myotubes, which may be clinically beneficial in the control of metabolic diseases.

## Introduction

Skeletal muscle is a major site of fatty acid oxidation (FAO) which has been shown to at least in part be regulated by the nuclear receptor transcription factor Peroxisome Proliferator Activated Receptor Delta (PPARδ)^1–4^. PPARδ activation is sensitive to nutrient availability in situations such as exercise^5^, fasting^6^ and high fat diets^7^ where there is an increase in circulating lipid concentrations. Indeed, poly-unsaturated fatty acids (PUFA) have been shown to bind effectively to PPARδ^8,9^ and therefore may have the potential, as natural ligands, to regulate fat oxidation in skeletal muscle, which may in turn prove beneficial for a number of diseases associated with excess lipid storage^10^.

In skeletal muscle cells the use of specific PPARδ agonists has elevated the expression of a number of proteins implicated in lipid entry, transport and catabolism^1,2^ and ultimately FAO^2^. PPARδ also regulates mitochondrial density and oxidative capacity^11,12^ which will in turn contribute to an increased capacity for FAO. Indeed, administration of the PPARδ agonist GW501516 prevents diet induced obesity and improves insulin sensitivity in genetically obese mice and those fed high fat diets^13^.

At present however, it is not known whether individual fatty acids, as natural PPARδ agonists, can influence lipid metabolism and mitochondrial biogenesis. Therefore here we aimed to determine whether selected PUFA which are known to bind efficiently to PPARδ^9^ can influence mRNA expression of a number of regulators of fat metabolism and mitochondrial density in cultures of primary human skeletal muscle cells. We found that the PUFA γ-Linolenic acid induced elevations in the expression of key regulators of lipid metabolism, and as such may provide a favourable dietary treatment in the attenuation of metabolic diseases.

## Methods

### Cell culture

Skeletal muscle biopsies were taken from the medial portion of the Vastus Lateralis of three healthy males aged 23.3 ± 0.9 years using the modified Bergstrom technique^14^. Ethical approval was obtained from Loughborough University Ethical Committee (R14-P64) and informed written consent was provided by all individuals before biopsies were taken. Primary human skeletal muscle derived cells (H-MDCs) were thereafter obtained from the biopsies using the explant technique as previously described^15^, and cells were maintained in growth media (GM) consisting of high glucose DMEM (Sigma Aldrich, Haverhill, UK) supplemented with 20% FBS (Pan Biotech, Aidenbach, Germany), 100U/ml penicillin and 100µg/ml streptomycin (GIBCO, Paisley, UK).

All experiments were conducted in 12 well culture plates (Fisher Scientific, Loughborough, UK), and for mitochondrial staining, 0.2% gelatin coated 13mm coverslips were placed in the wells. 12.5 × 10^3^ H-MDCs per cm^2^ were seeded into each well and were cultured in GM until confluent. Thereafter, GM was replaced with differentiation media (DM) consisting of high glucose DMEM supplemented with 2% Horse serum (Sigma), 100U/ml penicillin and 100µg/ml streptomycin (GIBCO) in order to encourage myotube formation. Following 72 hours in DM and myotube identification, H-MDCs were treated for a further 72 hours with DM supplemented with 1% fatty acid free BSA (Sigma) and either 10nM of the PPARδ agonist GW501516 (BioVision, Milpitas, USA), 250µM of Eicosapentaenoic acid (EPA, Cayman Chemicals, Ann Arbor, USA), 250µM γ-linolenic acid (γ-LA, Cayman Chemicals) or DM containing 6µl of ethanol to act as a vehicle (CON). Following treatment, cells were either stained with MitoTracker® Red CMXRos (Life Technologies, Warrington, UK) and subsequently fixed for fluorescence microscopy or lysed for RNA analysis. All experiments were conducted using cells between passage 3 and 6.

### Mitochondrial staining

Following 72 hours of treatment, cells were washed twice in PBS and then incubated for 45 minutes with 100nM MitoTracker® Red CMXRos in serum free media, and subsequently fixed with 3.7% paraformaldehyde. Cells were then permeablised with 0.2% triton X-100 diluted in TBS for 30 minutes and counterstained with DAPI prior to being mounted to glass microscope slides using Fluoromount™ (Sigma) mounting medium. From each of the mounted coverslips, 7 fluorescence images (Image size: x20, 641×479um) were taken at random using DM 2500 Fluorescence microscope (Leicia, Milton Keynes, UK). Mitochondrial density was quantified through measuring the mean stain intensity per myotube using the red channel only and maintaining constant exposure for each image. All image analysis was conducted on ImageJ analysis software (v1.48, National Institutes of Health, USA).

### RNA extraction and RT-PCR

At the end of the culture period, cells were washed twice in PBS and lysed in 500µl of TRIzol® (Life Technologies). RNA was thereafter extracted according to the manufacturer’s instructions and quantified using UV spectroscopy (Nanodrop, Thermo Fisher, Loughborough, UK). q-PCR reactions were prepared in 384 well plates, and consisted of 20ng of RNA reconstituted at 4ng/µl in 5µl of nuclease free water. 0.1μl of both forward and reverse primers (Table 1), 0.1μl of Quantifast Reverse Transcriptase kit (Qiagen, Crawley, UK) and 4.7μl of SYBR green mix (Qiagen) were added to the RNA to create 10μl reactions. q-PCR was performed using a ViiA 7™ Real Time PCR System (Applied Biosystems, Life Technologies, UK) programed to perform the following steps: 10min hold at 50°C (reverse transcription), followed by a 5min hold at 95°C (activation of ‘hot start’ Taq polymerase), and cycling between 95°C for 10s (denaturation) and 60°C for 30s (annealing and extension). Fluorescence was detected after every cycle and data was analysed using POLR2B as the housekeeping gene as a comparator to genes of interest (Table 1). Data was made relative using the comparative 2^−ΔΔCT^ (16) using the CON condition as a calibrator sample for each experimental repeat.

**Table 1.**
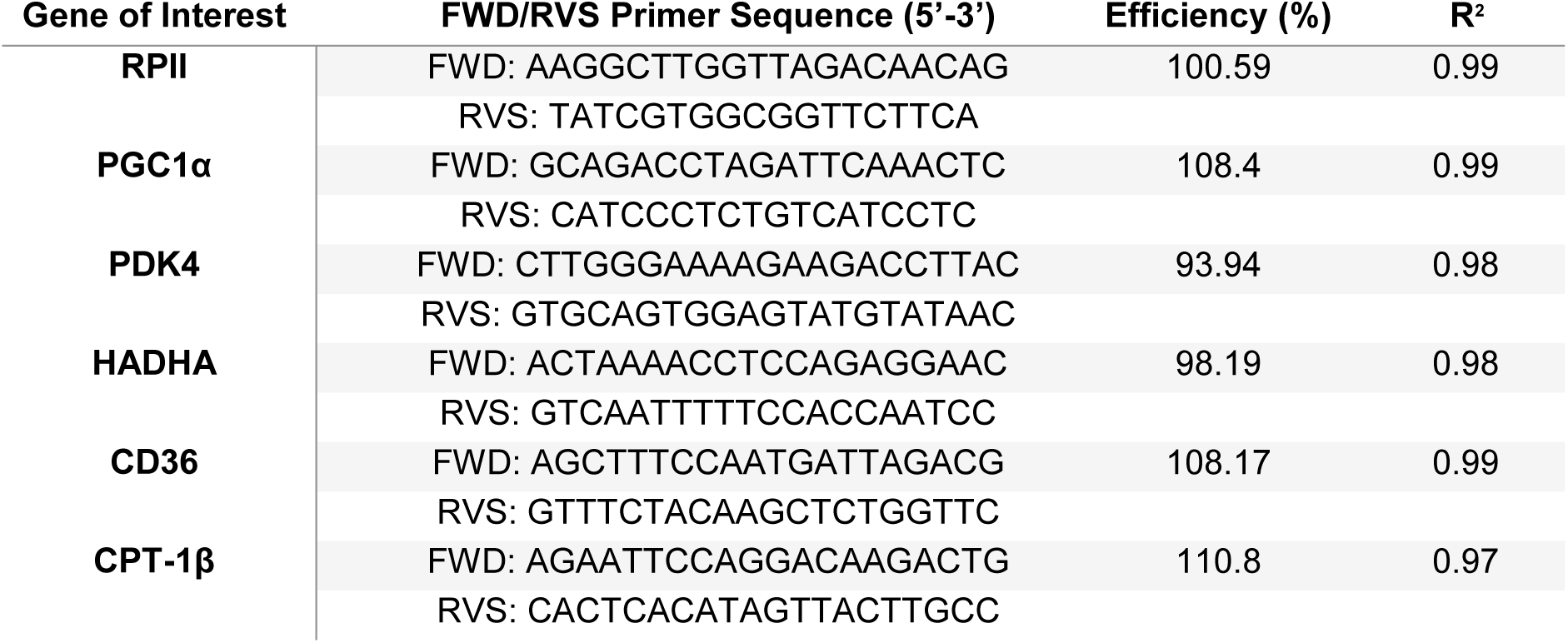
Primer table detailing forward and reverse primers utilised for the measurement of mRNA expression via q-PCR.

### Statistical Analysis

Data are presented as means ± SEM for 6 experimental repeats (from 3 separate muscle biopsies). Statistical analyses were performed using SPSS v.17 (SPSS Inc., Chicago, IL, US). Data was tested for normal distribution and homogeneity of variance. A one-way ANOVA or non-parametric equivalent was used to compare means across conditions for all variables and where significance was reported, Bonferroni post-hoc tests or non-parametric equivalent were used to detect where any significance lay between conditions. Effect sizes (r) were also used to interoperate the magnitude of effect as trivial (<0.2), small (0.2-0.5), moderate (0.5-0.8) or large (0.8≥).

## Results and Discussion

Incubation of human myotubes with γ-LA for 72 hours resulted in a significant increase in the mRNA expression of the long chain fatty acid transmembrane receptor CD36 compared with untreated myotubes (2.62 vs. 1.07, *U*= 23, z= −2.75, p= 0.005, r= 1.09). Interestingly however, neither EPA nor the PPARδ agonist GW501516 elicited an increase in CD36 expression. The mRNA expression of Carnitine Palmitoyltransferase (CPT1β); which catalyses the conversion of Fatty Acyl CoA to Acyl Carnitine, in turn allowing the activated fatty acid to enter the mitochondrial matrix to undergo β-Oxidation, was not different between any conditions (*F*_(3,47)_= 0.13, p= 0.945). In a manner similar to that shown for CD36, mRNA levels of the β-oxidation enzyme that catalyses the oxidation of straight-chain 3-hydroxyacyl-CoAs, β-HAD (HADHA) was elevated after incubation of myotubes with γ-LA (1.25 vs. 0.94, *F*_(3,52)_= 3.25, p= 0.022, r= 0.98) whereas no change was observed in the other conditions. Pyruvate dehydrogenase kinase (PDK4) is responsible for the phosphorylation of Pyruvate dehydrogenase, resulting in its inhibition and triggering a fuel utilisation switch to fatty acid metabolism. There was a pronounced increase in PDK4 mRNA expression in γ-LA treated cells, (2.90 vs. 1.25, *F*_(3,44)_= 3.04, p= 0.025, r= 1.67), and similar increases in PDK4 expression following treatment with EPA (2.47 vs. 1.25, *F*_(3,44)_= 3.04, p= 0.210, r= 1.51) and GW501516 (2.48 vs. 1.25, *F*_(3,44)_= 3.04, p= 0.570, r= 2.19), although the latter conditions did not reach statistical significance (Figure 1).

**Figure 1.**
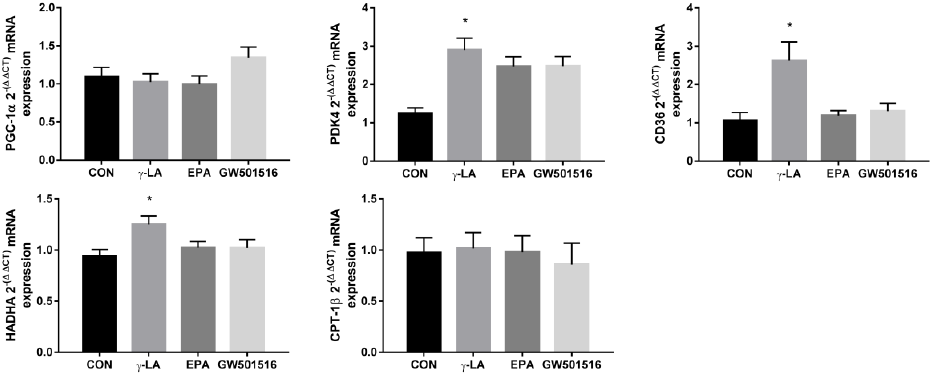
mRNA gene expression of PGC-1α, PDK4, CD36, HADHA and CPT-1β in human derived myotubes post 72h supplementation with one of four conditions: CON, γ-LA, EPA and GW501516. * = p ≤ 0.05 when comparing the indicated condition with the CON condition. n=6 across 3 biological repeats.

We also sought to determine whether these fatty acids could impact overall mitochondrial content by analysis of MitoTracker® red staining and PGC-1α mRNA levels. Interestingly we found that MitoTracker® staining intensity was greatest in γ-LA (36.81 vs. 25.59, *F*_(3,56)_= 7.80, p≤ 0.001, r= 1.45) and EPA treated myotubes (34.11 vs. 25.59, *F*_(3,56)_= 7.80, p= 0.005, r= 1.26); however there was no increase in PGC-1α expression in these conditions (Figure 1). Incubation of myotubes with GW501516 led to mean increases in both MitoTracker® staining intensity (33.03 vs. 25.59, *F*_(3,56)_= 7.80, p= 0.205, r= 1.52) and PGC-1α mRNA expression (1.34 vs. 1.09, *F*_(3,46)_= 0.73, p= 0.270, r= 0.58) compared to CON, although these increases did not reach statistical significance (Figure 2).

**Figure 2.**
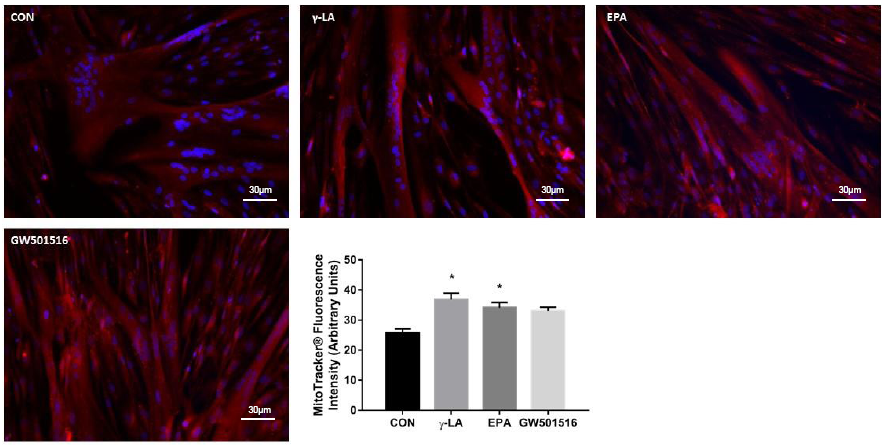
Mitotracker red staining as an indication of mitochondrial abundance was quantified in human derived myotubes post 72h incubation with one of four conditions: CON, γ-LA, EPA or GW501516. * = p ≤ 0.05. n=6 across 3 biological repeats.

In conclusion, these data provide the initial rational that the long chain unsaturated fatty acid γ-LA can induce the transcription of key regulators of lipid metabolism in human skeletal myotubes alongside increases in mitochondrial density. With a growing prevalence of diseases associated with obesity^10^, increasing FAO through dietary supplementation may provide a useful clinical aid, and warrants further investigation.

## Conflicts of Interest

The authors declare that there is no conflict of interest.

